# Spatial Variability in Streambed Microbial Community Structure Across Two Watersheds

**DOI:** 10.1101/2021.01.28.428737

**Authors:** Philips O. Akinwole, Jinjun Kan, Louis A. Kaplan, Robert H. Findlay

## Abstract

The spatial patterns of microbial communities are largely unknown compared to those of macro-fauna and flora. We investigated patterns of microbial community structure on streambed sediments from two watersheds across spatial scales spanning < 1m within a single stream to several hundred km between watersheds. Analyses of phospholipid fatty acids (PLFA) profiles indicated that the variations in microbial community structure were driven by increases in the relative abundance of microeukaryotic photoautotrophs and their contribution to total microbial biomass. Furthermore, streams within watersheds had similar microbial community structure, underscoring within-watershed controls of microbial communities. Moreover, bacterial community structure assayed as either polymerase chain reaction-denaturing gradient gelelectrophoresis (PCR-DGGE) fingerprints or PLFA profiles edited to remove microeukaryotes indicated a distinct watershed-level biogeography. No distinct stream order-level distributions were identified although DGGE analysis clearly indicated that there was greater variability in community structure among 1st-order streams compared to 2nd- and 3rd-order streams into which they flowed. Longitudinal gradients in microbial biomass and structure showed that the greatest variations were associated with 1st order streams within a watershed and 68% of the variation in total microbial biomass was explained by sediment C:N mass ratio, percent Carbon, sediment surface area, and percent water content. This study confirms a distinct microbial biogeography for headwater stream communities driven by environmental heterogeneity across distant watersheds and suggests that eukaryotic photoautotrophs play a key role in structuring sediment microbial communities.

**IMPORTANCE:** Microorganisms in streams drive many biogeochemical reactions of global significance, including nutrient cycling and energy flow, yet the mechanisms responsible for the distribution and composition of streambed microbial communities are not well known. We sampled sediments from multiple streams in two watersheds; Neversink River (New York) and White Clay Creek (Pennsylvania) watersheds and measured microbial biomass, total microbial and bacterial community structures using phospholipid and molecular methods. Microbial and bacterial community structures displayed a distinct watershed-level biogeography. The smallest headwater streams within a watershed showed the greatest variation in microbial biomass, and C:N ratio, percent carbon, sediment surface area and percent water content explained 68% of the variations in microbial biomass. This study indicates a non-random distribution of microbial communities in streambeds, and that microeukaryotic photoautotrophs, environmental heterogeneity and geographical distance influence microbial composition and spatial distribution.

## INTRODUCTION

Microorganisms are the most biologically diverse and ubiquitous taxa on earth and their metabolic activities largely control biogeochemical cycling and many ecosystem processes (1–3). In stream ecosystems, benthic microbial communities mediate biochemical transformations, including degradation and transformation of organic compounds into biomass or inorganic components, and exert significant control over the mineralization and downstream export of terrestrially-derived dissolved organic matter (DOM) (4–8). In addition, microbial processing of terrestrial DOM and nutrients within streambed sediments is essential to material flux to higher trophic levels (5, 9-11). However, microorganisms are often ignored or highly aggregated in many stream food web analyses (12).

While the central role of microorganisms in ecosystem functions is well documented, the linkage between microbial community composition and ecosystem functions remains elusive (13), with streams being among the least studied ecosystems. It has been suggested that streams function as meta-ecosystems (14) with a longitudinal acclimation of microbial communities as streams get larger (i.e., increasing stream order) within a drainage network (15). The focus on a longitudinal ecological framework emphasizes central differences among streams and lentic aquatic habitats and underpins a suite of conceptual and mathematical models: River Continuum Concept (16); Flood Pulse Concept (17, 18); Pulse Shunt Hypothesis (19); and nutrient spiraling (20) used to guide stream ecosystem research. These differences include association of downstream movement with organisms, energy flow, and nutrient cycling within a physical network where strong directionality and alteration to fluvial geomorphology and hydrology influence energy sources and biogeochemical processes (21). Combined, these characteristics require spatially explicit sampling over a range of scales to better understand key physicochemical and biological processes (21). For example, fine-scale sampling and spatially specific analysis have shown that chemical constituents in stream water exhibit either flow-connected, flow-unconnected, or Euclidean spatial relationships (22).

The existence of biogeographic patterns that span spatial scales over seven orders of magnitude have been established for a wide range of microorganisms (23), and although the mechanisms shaping these patterns have not been identified (24), the drivers of microbial diversity clearly depends upon spatial scale (25, 26). The species-area and the distance-decay relationships commonly observed for macroscopic organisms are also common for microbial communities although spore formation or dormant vegetative stages among microbes may contribute to species persistence and reduce the rate of species turnover for microbial communities (27). Relatively few studies have addressed temporal or spatial patterns of heterotrophic microbial community composition in streams, but a clear annual recurrence of taxa within a single stream (28) and a biome-level pattern in microbial community structure for streambed sediments have been observed (29). Alternatively, a study of nine streams across the southeastern and midwestern United States attributed differences in microbial community structure to variations in chemical characteristics of the habitats, rather than a pattern driven by spatial gradients (30). Factors contributing to the structure and function of streambed microbial communities include: sunlight and water flow (31); water temperature and desiccation (32); hydrology (33–35); pH, sediment grain size, inorganic nutrients, and dissolved oxygen (5); bedrock type (36); inter-specific competition, viral lysis and flagellate grazing (37); anthropogenic pollution (38, 39); and DOM concentration and quality (36, 40). For instance, the dominant controls on benthic microbial diversity such as sunlight and algal abundance contributes to the relative importance of algal versus terrestrial-derived DOM substrates in lotic systems (41). Changes in light levels have been shown to affect phototrophic growth rates (42). However, the spatial scales at which these local environmental gradients give way to biogeographical processes as the major determinants of microbial community structure and biogeochemical functions have yet to be determined (8).

Given the potential for hydrologic disturbance and metacommunity dynamics to determine microbial community structure in fluvial networks (43) and the higher alpha and beta diversities reported for biofilms growing on rock surfaces in headwater streams compared to higher-order streams within a fluvial network (44), we hypothesize that microbial communities on streambed sediments should exhibit spatially complex, but longitudinally distinct patterns. More specifically, if patterns observed for biofilms on rocks (44) hold for biofilms on the more easily disturbed substratum of streambed sediments, variability in bacterial and microbial community structure among headwater streams should be greater than that found within larger streams of the same network.

To test this hypothesis, we examined microbial community structure and biomass from streambed sediments in low order streams within two eastern deciduous forest watersheds of the Delaware River - White Clay Creek within the Pennsylvania Piedmont and Neversink River in the Catskill Mountains of New York. We sampled using a nested design that spanned five orders of magnitude across four spatial scales (Fig. 1). We analyzed microbial phospholipid fatty acids (PLFA) to assess total microbial community structure at the resolution of functional groups, performed molecular methods (i.e. PCR-DGGE) to assess individual taxa, and used phospholipid phosphate-based analyses to quantify total microbial biomass. Multivariate statistics were used to compare total microbial biomass, microbial and bacterial community structure across distant watersheds.

**Figure 1.**
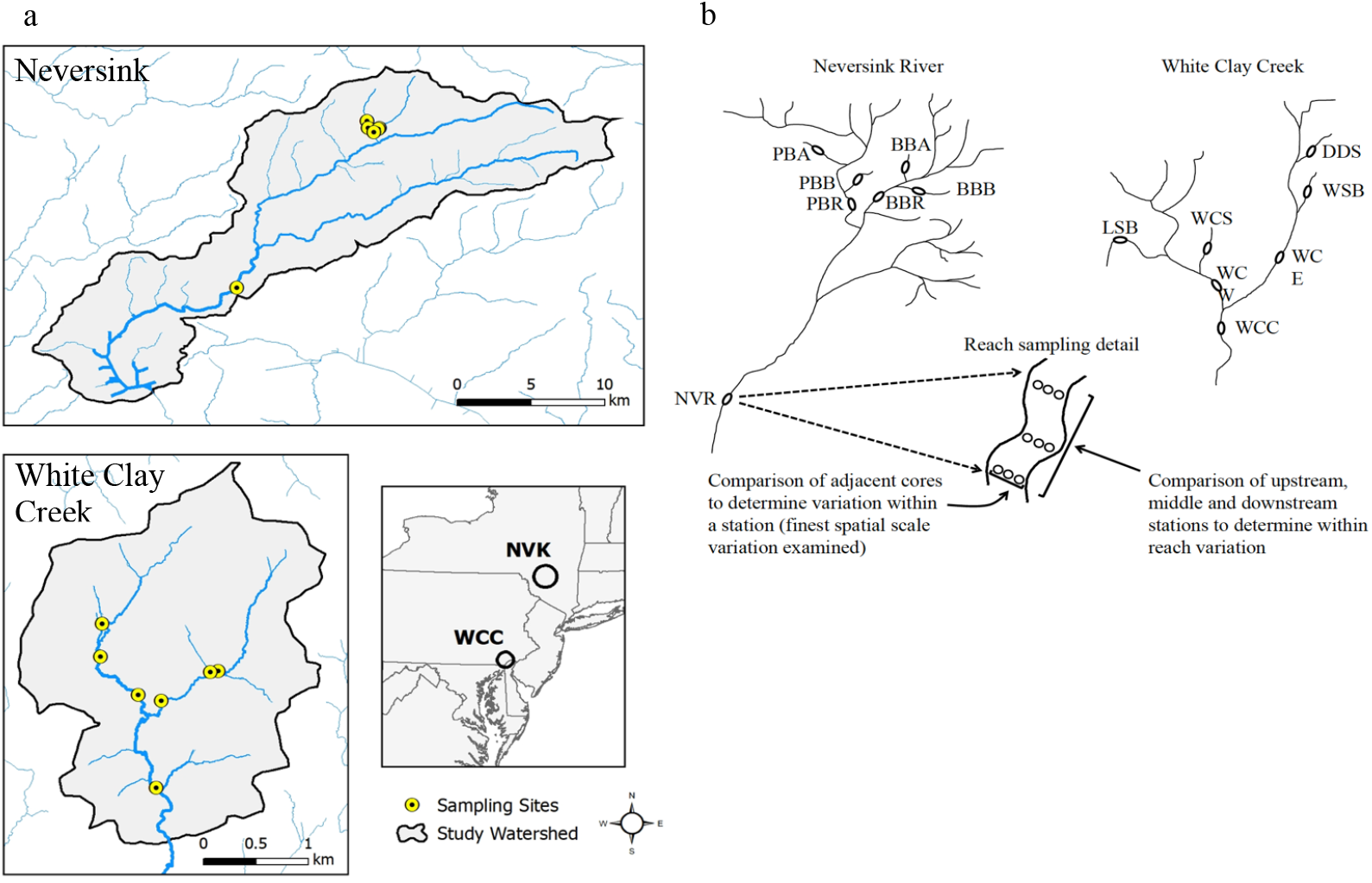
(a) Map of the Neversink watershed in New York and White Clay Creek watershed in Pennsylvania, USA (b) Sampling scheme used to examine microbial biomass and community structure across multiple spatial scales in the two watersheds. Sampling within the Neversink watershed consists of four 1^st^ order streams; Biscuit Brook and Pigeon Creek tributaries (Biscuit Brook Tributary A and B [BBA, BBB], Pigeon Creek Tributary A and B [PBA, PBB]), two 3^rd^ order streams (Biscuit Brook [BBR] and Pigeon Creek [PBR]) and one 5^th^ order stream (Neversink River [NVR]). Sampling within the White Clay Creek watershed consists of four 1^st^ order streams (Ledyards Spring Branch [LSB], Water Cress Spring [WCS], Dirty Dog Spring [DDS] and Walton Spring Branch [WSB]), two 2^nd^ order streams (East and West Branch White Clay Creek [WCE, WCW, respectively]) and one 3^rd^ order stream (White Clay Creek [WCC]). Sketches of watersheds are not drawn to scale. Each eclipse represents a reach, which contained 3 stations, each of which was sampled 3 times.

## RESULTS

### Microbial community structure

Across both watersheds, the major component of variation in streambed microbial community structure was the relative contribution of bacteria and microeukaryotes to these communities. Fatty acids indicative of microeukaryotes (20:4ω6, 18:2ω6), photosystem II (16:1ω13t), and chrysophytes and chlorophytes (18:3ω3, 20:5ω3, 16:3ω4) (45) were present in greater relative abundances for streams with more negative principal component 1 (PC1) loadings while the bacterial fatty acids (cy17:0, cy19:0, a17:0, i17:0, i15:0, br17:1a and 10me16:0) were present in greater relative abundances for those streams with more positive PC1 loadings (Fig. 2). Within the Neversink River watershed the contribution of microeukaryotes ranged from 42% (NVR) to 12% (PBB), and in the White Clay Creek watershed the range was 31% (WSB) to 4% (DDS) (Table 1). The percentage that microeukaryotes comprise of total microbial biomass and PC1 factor scores showed a strong, negative correlation (r^2^ = 0.88; Fig. 3a). Thus, the position of stream communities along PC1 were arrayed according to the proportions of prokaryotes and microeukaryotes within communities, moving from negative to positive PC 1 loadings as the relative contribution of microeukaryotes decreased. PC1 separated the two streams (NVR and WSB) that showed the highest relative contribution of microeukaryotes from all other streams (Fig. 2). To further investigate the role of microeukaryotic community structure on this relationship, percent contribution of microeukaryotes to total biomass was compared to the ratio of fatty acid markers for phototrophs (ω3) to fatty acid markers for heterotrophs (ω6). The strong positive linear correlation between these two parameters indicated that increasing importance of microeukaryotes within the microbial community correlated with increasing importance of phototrophs within the microeukaryotic community (r^2^ = 0.71; Fig. 3b). Furthermore, in figure 2 the streams (excepting NVR) separated by watershed along principal component 2 (PC2) as WCC streams had positive PC2 scores while the Neversink streams had negative PC2 scores. Within a watershed, microbial community composition of streambed sediments from 1^st^-, 2^nd^- and 3^rd^-order streams were, in general, similar although three 1st-order streams (WSB, WCS and DDS) in the White Clay Creek watershed separated from the two 2nd-order streams (WCW and WCE) (Fig. 2).

**Figure 2.**
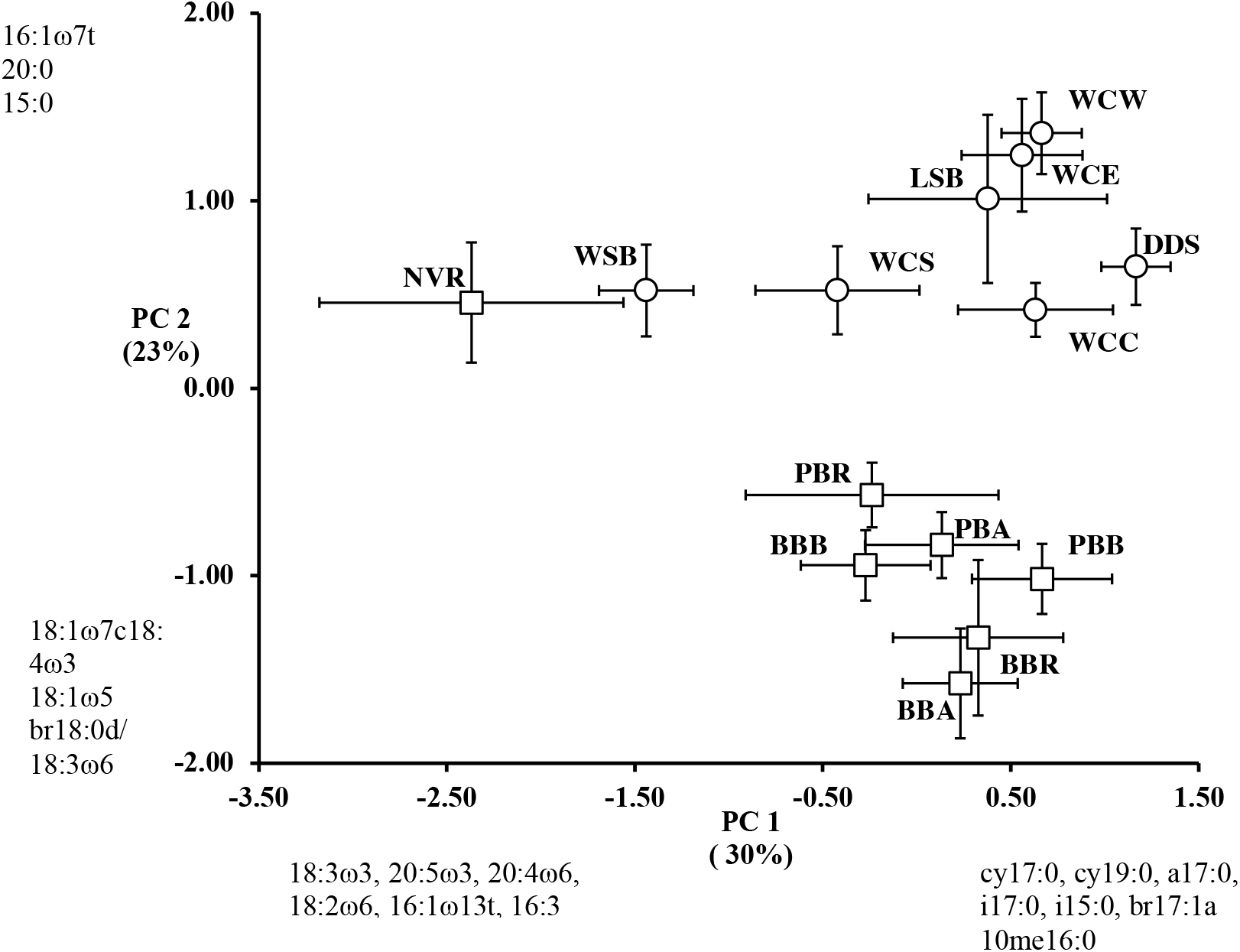
Principle Component Analysis of stream sedimentary microbial community structure of PLFA profiles of White Clay Creek (open circle) and Neversink (open square) watersheds. The percent variation explained by each axis is indicated on the respective component axis. Identified fatty acids had component loadings of >|0.5| with strong influence on the pattern of variation among samples along the respective component axes. Site abbreviations are as described in the legend to Fig.1

**Figure 3.**
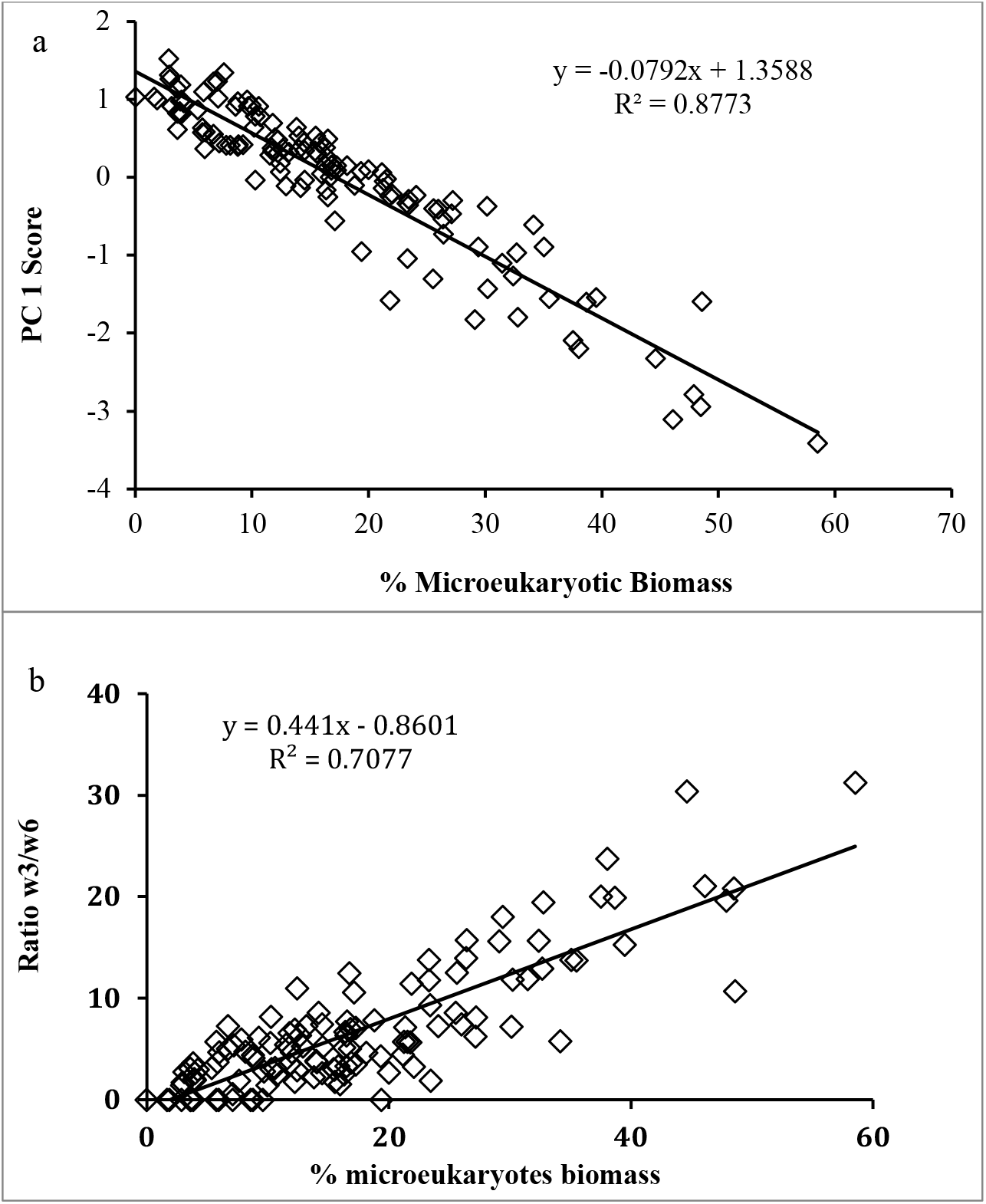
a) Relationship between Principle Component Analysis factor 1 score and the calculated percentage that microeukaryotes contribute to total microbial biomass for all stream samples, b) Relationship between the calculated percentage that microeukaryotes contribute to total microbial biomass and the ratio of ω3 to ω6 fatty acids for all stream samples from the PLFA profiles.

**Table 1.** Microbial biomass and composition, water chemistry and sediment organic content of White Clay Creek and Neversink watersheds.

| Watershed<br>& stream | Biomass/PLP<br>(nmol g<sup>-1</sup> fresh<br>wet wt)<sup>a</sup> | Bacterial<br>abundance<br>(g<sup>-1</sup> fww)<sup>b</sup> | %<br>Eukaryotic/<br>prokaryotic<sup>c</sup> | Cond<br>(μS/cm) | Temp<br>(°C) | % C | % N | C:N ratio<sup>d</sup> |
| --- | --- | --- | --- | --- | --- | --- | --- | --- |
| White Clay Creek |  |  |  |  |  |  |  |  |
| DDS | 14.80 ±2.81 | 5.67 x 10<sup>8</sup> | 4/96 | 186.0 | ND | 0.33 ±0.01 | 0.02 ±0.55 | 14.14 ±3.18 |
| WSB | 30.34 ±9.81 | 8.24 x 10<sup>8</sup> | 31/69 | 225.0 | ND | 0.58 ±0.02 | 0.04 ±0.54 | 12.98 ±2.53 |
| LSB | 43.29 ±19.60 | 1.56 x 10<sup>9</sup> | 9/91 | 373.6 | ND | 2.96 ±0.37 | 0.31 ±0.51 | 12.77 ±5.12 |
| WCS | 52.41 ±4.87 | 1.68 x 10<sup>9</sup> | 20/80 | 303.3 | ND | 3.36 ±0.15 | 0.21 ±0.34 | 16.45 ±2.09 |
| WCE | 22.13 ±3.38 | 8.26 x 10<sup>8</sup> | 6/94 | 189.9 | ND | 1.15 ±0.04 | 0.07 ±0.37 | 16.27 ±2.75 |
| WCW | 26.89 ±2.52 | 1.01 x 10<sup>9</sup> | 6/94 | 259.7 | 16.2 | 1.73 ±0.11 | 0.11 ±0.37 | 16.59 ±2.71 |
| WCC | 12.12 ±4.95 | 4.25 x 10<sup>8</sup> | 11/89 | 232.7 | 16.1 | 0.57 ±0.04 | 0.06 ±5.03 | 9.08 ±2.96 |
| Neversink |  |  |  |  |  |  |  |  |
| BBA | 6.99 ±2.39 | 2.27 x 10<sup>8</sup> | 19/81 | 35.1 | 15.4 | 0.36 ±0.01 | 0.03 ±1.08 | 11.15 ±2.30 |
| BBB | 10.95 ±3.74 | 3.34 x 10<sup>8</sup> | 23/77 | 32.4 | 14.8 | 0.34 ±0.01 | 0.03 ±0.55 | 10.25 ±3.15 |
| PBA | 12.18 ±3.78 | 3.92 x 10<sup>8</sup> | 18/82 | 25.6 | 16.2 | 0.72 ±0.04 | 0.05 ±0.35 | 12.99 ±3.10 |
| PBB | 16.42 ±6.96 | 5.73 x 10<sup>8</sup> | 12/88 | 18.8 | 15.5 | 1.18 ±0.11 | 0.08 ±0.38 | 11.06 ±3.30 |
| BBR | 8.54 ±1.79 | 2.87 x 10<sup>8</sup> | 16/84 | 20.8 | 16.4 | 0.18 ±0.00 | 0.03 ±0.30 | 5.62 ±2.38 |
| PBR | 6.77 ±0.75 | 2.03 x 10<sup>8</sup> | 25/75 | 24.5 | 16 | 0.16 ±0.01 | 0.02 ±0.88 | 6.18 ±3.76 |
| NVR | 13.58 ±4.41 | 2.95 x 10<sup>8</sup> | 42/58 | 33.5 | 15.3 | 0.20 ±0.01 | 0.03 ±0.71 | 7.69 ±1.05 |

<sup>a</sup> Mean ± standard deviation (n = 9).
<sup>b</sup> Calculated from PLP x % prokaryotic (expressed as decimal fraction) and a conversion factor
of 100 nmol PLP = 4 x 10<sup>9</sup> cells
<sup>c</sup> Percentage that microeukaryotic contributes of total microbial biomass, calculated from PLFA
profiles (n = 9).
<sup>d</sup> Percent that C and N contribute to total sediment elementary atoms (n = 9)
ND= measurements were not taken

PCA analysis for within streams comparisons showed that microbial community composition displayed no consistent longitudinal variation among stations within a stream (Fig. 4), that is, there were no consistent differences among cores from the upper, middle and lower station of a reach. Rather for some stations the three replicate cores showed nearly identical microbial community structure (dashed circle Fig. 4a and b), while for other stations the three replicate cores differed greatly in microbial community structure (dotted ellipse Fig. 4a and b). In addition, cores from different stations within a stream (horizontal arrows Fig. 4), and in a few cases, cores from different streams within a watershed (vertical arrows Fig. 4) showed nearly identical microbial community structure.

**Figure 4.**
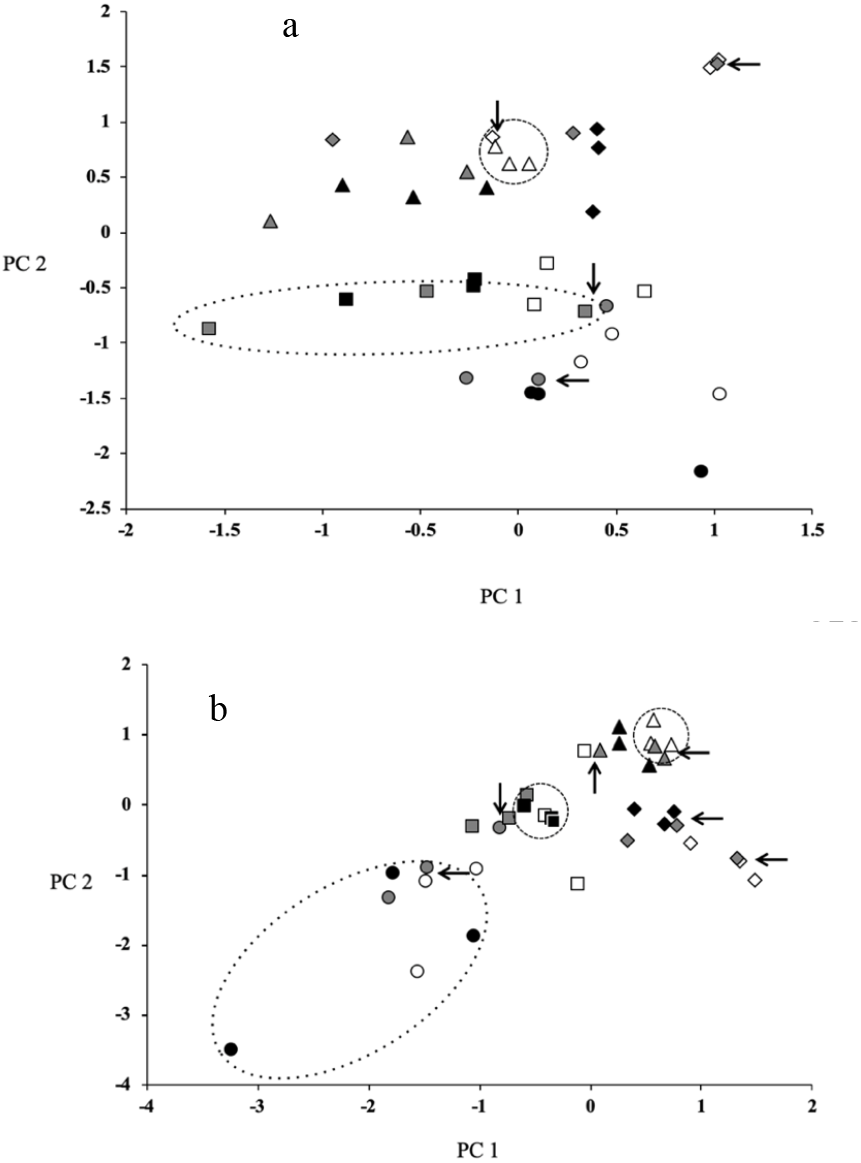
a) Comparison of microbial community structure among individual cores from four selected streams: Diamonds - LSB; triangles - WCS; squares - PBR; circles - BBR. Symbol color denotes station with white indicating upper reach, gray middle reach and black lower reach, respectively. The dashed circle indicates the station with lowest among adjacent core variation while the dotted ellipse indicates the station with the highest among adjacent core variation. The horizontal arrows indicate cores from different stations within a stream with nearly identical community structure while vertical arrows indicate cores from different streams with very similar community structure. b) Comparison of bacterial community structure among individual cores from four selected streams. Samples and their relationships are indicated as above with the addition of an upward pointing arrow indicating two samples from the two different watersheds with very similar community structure.

### Bacterial community structure

When bacteria alone are considered, community structure of streambed sediments separated out by watershed and this was consistent across the lipid and molecular approaches to assessing community structure; all streams in the Neversink River watershed had negative factor scores and all streams within the White Clay Creek watershed had positive factor scores (Fig. 5a and 5b). At the spatial scale of streams within a watershed, no clear longitudinal separation of stream orders was observed, although the variation in bacterial community structure among 1^st^ order steams assayed by DGGE was more variable than that observed for 2^nd^- and 3^rd^-order streams (fine vs. course dashed boxes Fig. 5b). At the spatial scale of replicate cores within a station (<1m), bacterial community structure assayed by PLFA showed patterns similar to those observed for microbial community structure (Fig. 4a and b).

**Figure 5.**
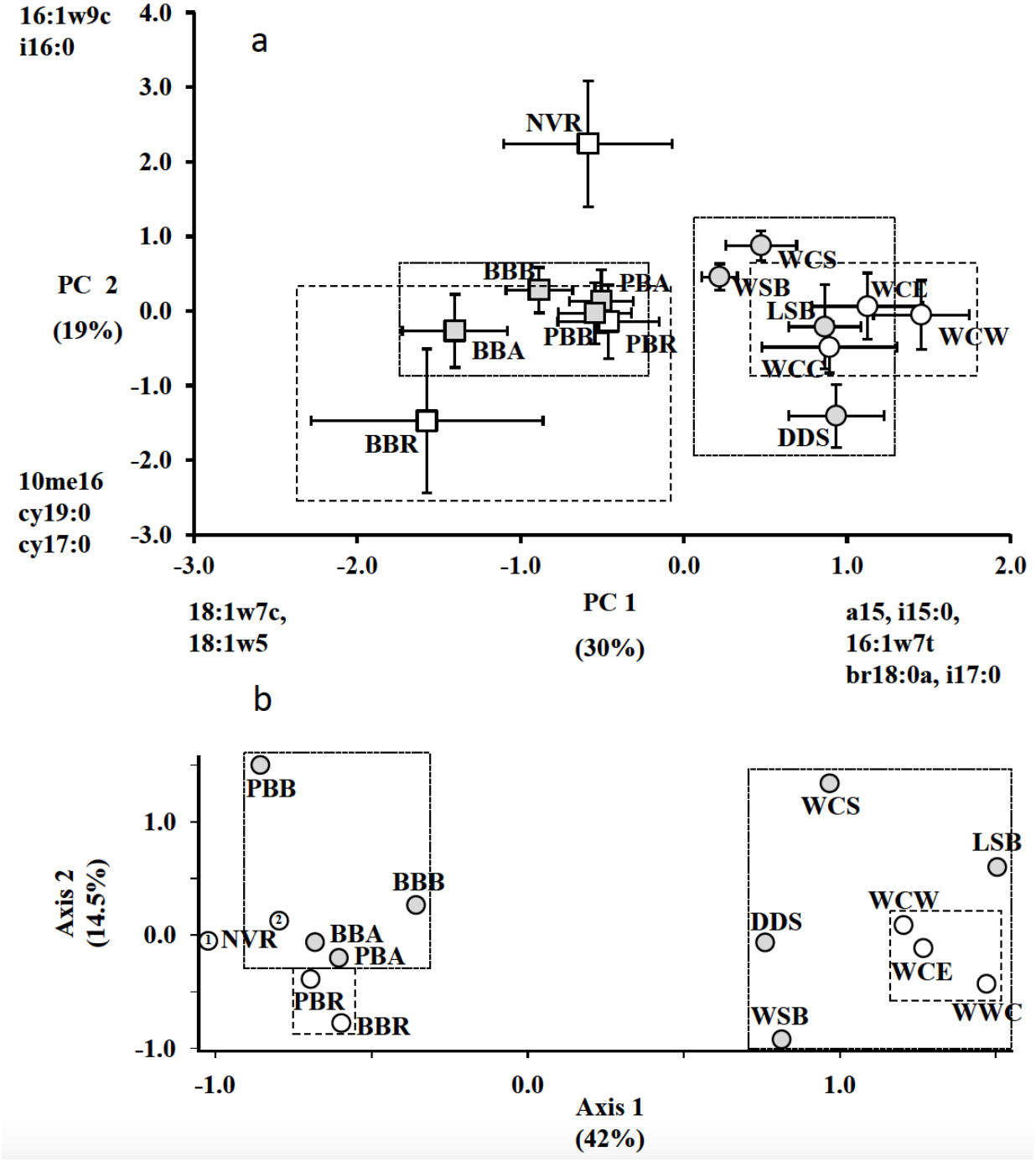
Spatial variation in sedimentary bacterial community composition in WCC and NVR watersheds. a) PCA analysis of PLFA profiles after removal of fatty acids assigned a priori to the functional group microeukaryotes and those known to be common to both bacteria and microeukaryotes from the PLFA profiles. b) NMDS analysis DGGE presence/absence data. Streams within the White Clay Creek dendritic network are denoted with a circle and those within the Neversink dendritic network with a square. Site abbreviations are as described in the legend to Fig.1. Dotted boxes indicate the relative variation in bacterial community structure of 1^st^ order stream sediments within a watershed and dashed boxes indicate the same for the 2^nd^ and 3^rd^ order streams or 3^rd^ order streams within the watershed.

### Total microbial biomass

Streams from the WCC watershed showed a greater range and a higher basin-wide average for total microbial biomass concentration, percent prokaryotic content, and bacterial abundance than streams within the Neversink River watershed (Table 1; Fig. 6). The microbial biomass differences were significant at the levels of watersheds (F= 15.18, p ≤ 0.005) and streams within watersheds (F= 7.38, p ≤ 0.001), but not stations within streams (Table 2, Fig. 6). At the scale of stations within streams (1m-50m), variability in sediment microbial biomass, expressed as a coefficient of variation, ranged from 30% to 79% with no consistent longitudinal pattern of biomass changes among stations (Fig. 6b). The coefficient of variation for microbial biomass among replicate cores within stations ranged from 5% to 83% (Fig. 6c).

**Figure 6.**
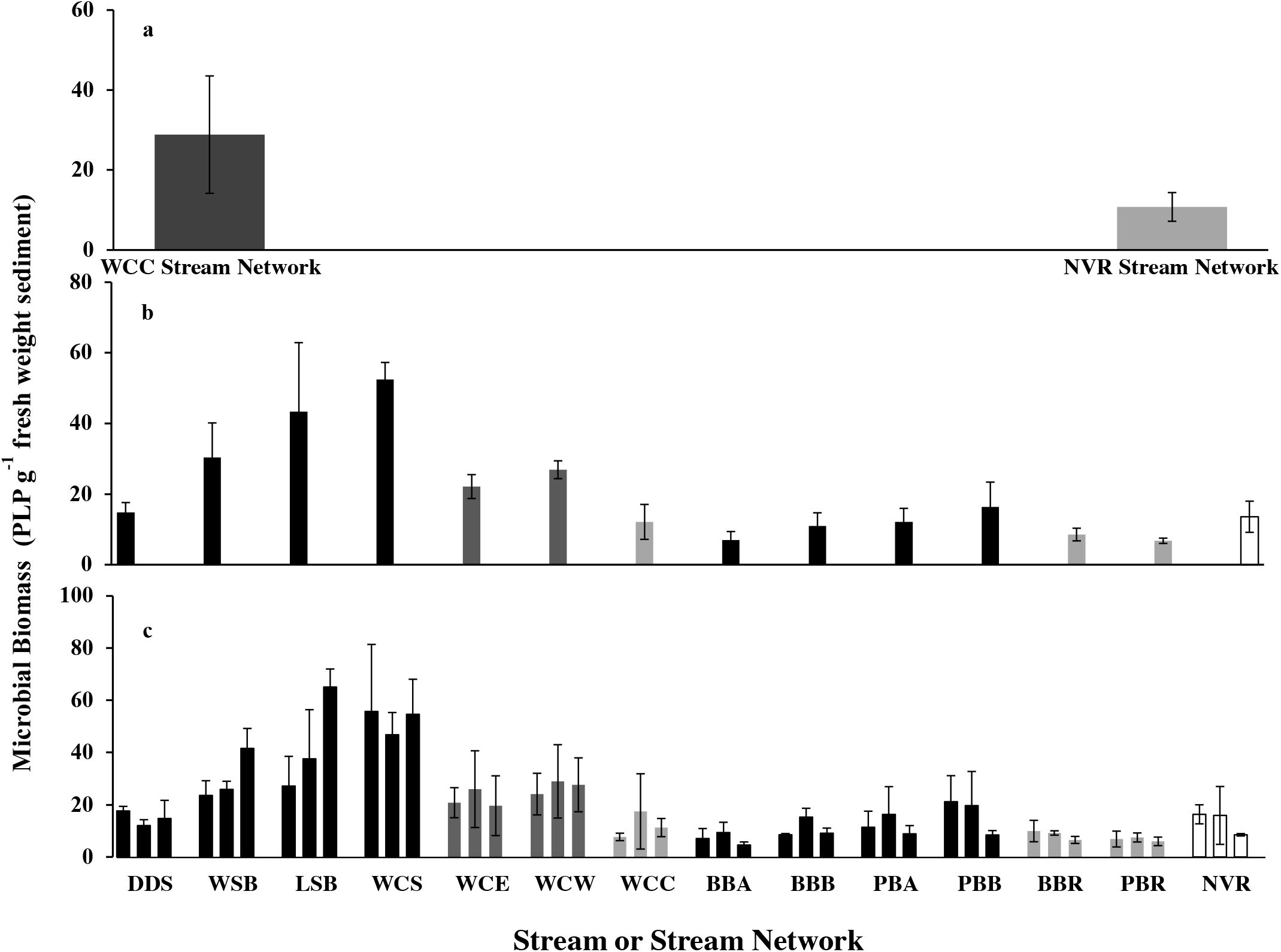
Microbial biomass (mean ± SD) of White Clay Creek and Neversink watershed sediments at three spatial scales: a; watershed, b; stream and c; station. Stream order (or average order for watershed values) are indicated as: black = 1st order, dark gray = 2nd order, light gray = 3rd order, open = 5 th order.

**Table 2.** Nested ANOVA to test the effects of watershed, streams within watershed, and stations within streams on microbial biomass.

| Source | DF | Adj SS | Adj MS | F | P |
| --- | --- | --- | --- | --- | --- |
| Watershed | 1 | 5.119 | 5.119 | 15.18 | 0.002 |
| Stream (Watershed) | 12 | 4.047 | 0.337 | 7.38 | 0.000 |
| Stations (Watershed*Stream) | 28 | 1.280 | 0.046 | 1.46 | 0.095 |
| Error | 84 | 2.628 | 0.031 |  |  |

### Water chemistry and sediment organic content

Sediment C and N content in streams from the two watersheds had overlapping ranges, but on average, sediments from the WCC watershed had 3.5x higher % C and 3x higher % N than the sediments from the Neversink watershed, and the C:N ratio was 1.5x higher in the WCC watershed (Table 1, Fig. 7). Streams from the WCW sub-basin in the White Clay Creek that has more carbonate bedrock, WCW and its 1st order tributaries, WCS and LSB, had greater C and N content compared to many other streams and few significant differences among the other streams (Fig. 7 a, b). In general, sediment C:N mass ratios were higher in 1^st^ and 2^nd^ order streams and lower in 3^rd^ and 5^th^ order stream sediments (Fig. 7c). Conductivity of stream water in the WCC watershed was highest in the WCC tributaries and ~ 5-fold greater than the conductivity of the stream water within the Neversink watershed (Table 1). Combined sediment percent carbon content, percent water content, C:N mass ratio and sediment surface area explained approximately two-thirds of the variation observed in sedimentary microbial biomass (Table 3, Model 7). Path analysis indicated that the variables percent carbon content, percent water content, and C:N ratio had significant direct effects on biomass and that sediment surface area was correlated, to a greater or lesser extent, with these three variables (Fig. 8). Two models were investigated to discern the theoretical linkage and directionality among the variables, one constrained and one unconstrained. The constrained model links sediment surface area indirectly to biomass via its direct effects on sediment carbon and water content, while the unconstrained model links surface area indirectly to biomass via its correlations with sediment carbon content, water content and C:N ratio. These two models yielded very similar results and we present only the unconstrained model. Percent carbon content showed the greatest direct effect on biomass (r^2^ = 0.393) as well as substantial indirect effects via its correlation with percent water content and C:N mass ratio (Fig. 8). Combined, the direct and indirect effects of carbon accounted for ~ 61% of the variation in total sediment microbial biomass. Similarly, percent water content and C:N mass ratio accounted for 56% and 37%, respectively, of the variation in total sediment microbial biomass. Sediment surface area via indirect effects accounted for ~34% of the variation in total sediment microbial biomass (Fig. 8).

**Figure 7.**
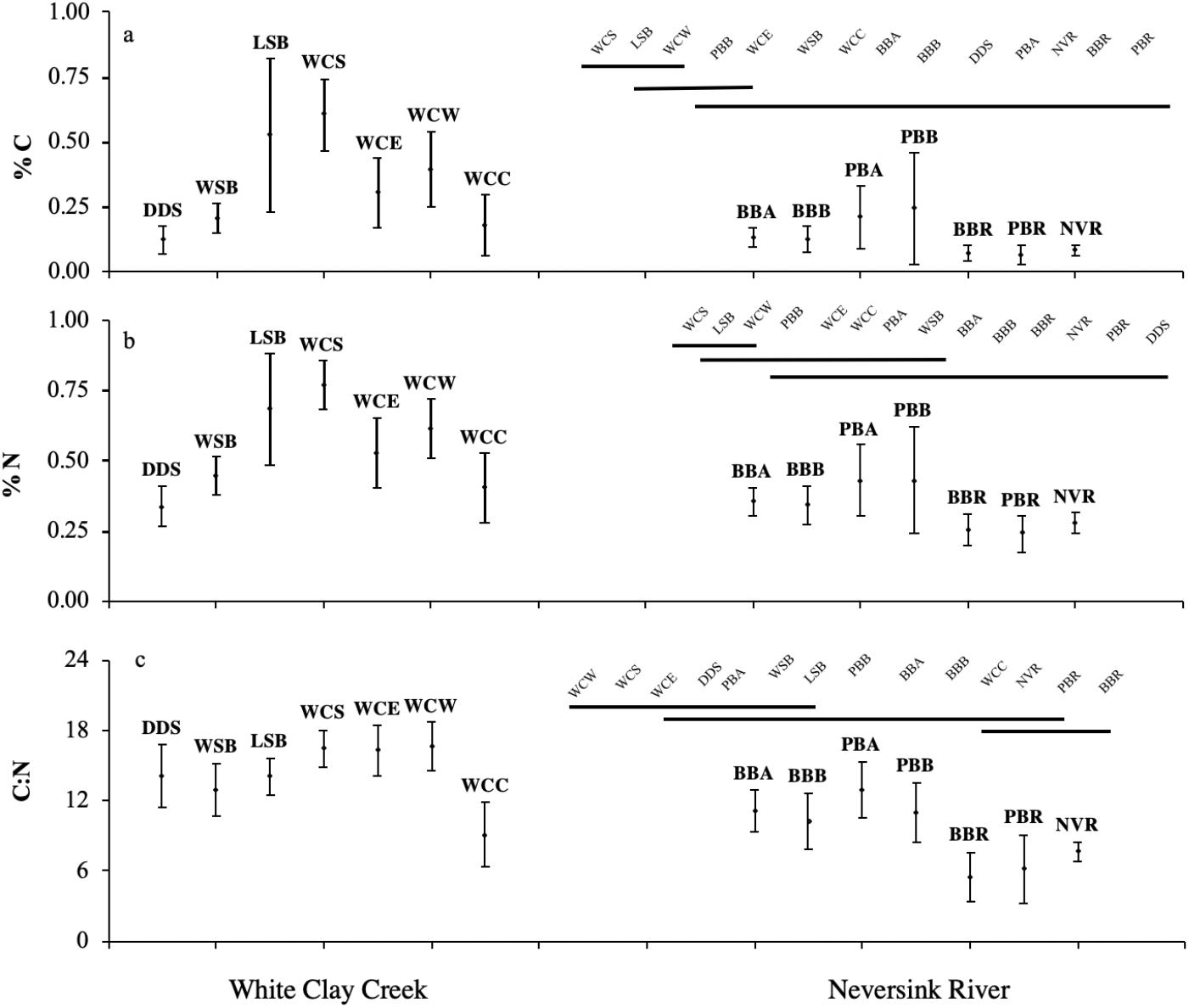
Variation in sediment (a) percent carbon, (b) percent nitrogen and (c) C:N mass ratio by stream order (1^st^ to 3^rd^ /5^th^ order from left to right) and watershed. Vertical bars denote 0.95 confidence intervals. Streams not connected by a horizontal line are significantly different (p = 0.05, Tukey’s Wholly Significant Difference).

**Figure 8.**
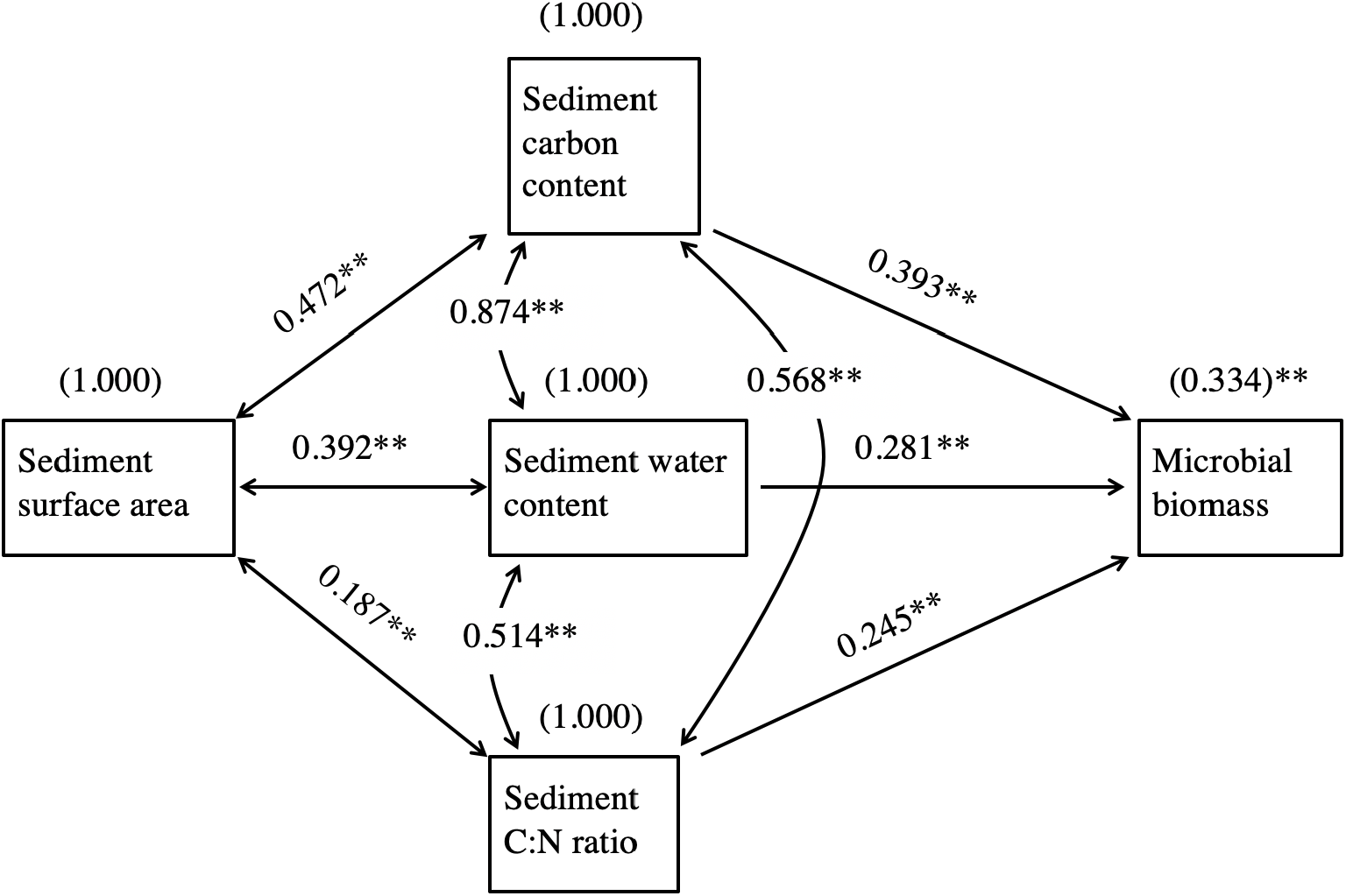
Path diagrams describing the structure of the relationship between sediment microbial biomass and % Carbon (%C), % water content, C:N mass ratios (C:N) and sediment surface area (SSA). Single-headed arrows indicate casual paths; numbers on arrows are path coefficients (standardized regression coefficients) indicating the relative strength of each path leading to a given response variable. Double-headed arrows represent the correlations among the predictor variables. Arrows connecting environmental variables to the independent variable (microbial biomass) indicate direct effects, while environmental variables linked to the independent variable via other environmental variable constitute indirect effects. Path coefficients calculated by SAS Structural Equation Modeling for JMP 10. *= P <0.01, **=P <0.001

**Table 3.** Multiple regression analysis (best subsets) for natural log biomass as a function of various physical and chemical stream parameters.

| Model | Vars | R-Sq(adj) | Mallows Cp* | SE | %Water | SSA | %C | %N | C:N |
| --- | --- | --- | --- | --- | --- | --- | --- | --- | --- |
| 1 | 1 | 60.4 | 30.2 | 0.2052 |  |  | X |  |  |
| 2 | 1 | 56.0 | 46.3 | 0.2161 | X |  |  |  |  |
| 3 | 2 | 64.4 | 15.9 | 0.1943 |  |  | X |  | X |
| 4 | 2 | 64.0 | 17.6 | 0.1955 |  |  | X | X |  |
| 5 | 3 | 66.7 | 8.4 | 0.1879 |  | X | X |  | X |
| 6 | 3 | 66.0 | 11.1 | 0.1899 | X |  | X |  | X |
| 7 | 4 | **68.1** | **4.3** | **0.1839** | **X** | **X** | **X** |  | **X** |
| 8 | 4 | 67.0 | 8.5 | 0.1872 | X | X | X | X |  |
| 9 | 5 | 68.0 | 6.0 | 0.1845 | X | X | X | X | X |

\*the model giving the minimum Mallows Cp statistic was used; Mallows (91).

## DISCUSSION

### Spatial variability of microbial community structure, and the role of phototrophic microeukaryotes

Our results indicate that microbial community structure in the fourteen headwater streams investigated within the Eastern Deciduous Forest displayed a distinct regional-level spatial variability, even when streams within a watershed displayed high within-stream or among-stream variations. Sediments from the WCC and Neversink River watersheds, with the exception of sediments from the 5th-order Neversink River site, differed in total microbial community structure; this difference was detected by the second component of variation. These findings extend previous studies that indicate geographical distance is important in structuring microbial communities at regional scales (26, 28, 29), and provide further evidence for spatial variations in microbial community structure (27). For instance, in a recent study, Zhang et al. (26) investigated soil microbial communities along a ca. 878-km transect during two contrasting seasons, and reported that spatial heterogeneity rather than seasonality explained more of the spatiotemporal variation of soil microbial alpha and beta diversities. In the Findlay et al. study (29) when sediment microbial community structure was compared among three biomes (eastern deciduous, south-eastern coniferous and tropical evergreen forests), microbial community within a biome were more similar in composition than community from different biomes that differed in environmental heterogeneity. A major difference between our study and that of Findlay et al. (29) was that the two watersheds examined in this study occurred within the same biome (Eastern Deciduous Forest). However, as documented in the study site descriptions, below, only one of the watersheds (Neversink) was glaciated during the last glacial period and the watersheds differ in soil age and structure, bedrock geology, and land cover. This study also showed that the WCC and Neversink watersheds differed in C:N mass ratio, % carbon, sediment surface area, and microeukaryotic/ prokaryotic ratio, suggesting that these factors, along with others, affect spatial variability of freshwater sediment microbial communities at a variety of scales within watersheds.

One major component of variation in microbial community structure reflects the ratio of microeukaryotic to prokaryotic biomass. As streams with high microeukaryotic biomass were dominated by phototrophs, and light has long been known as a major factor influencing the relative abundance of microeukaryotes within microbial communities (41, 46), canopy cover likely influenced the placement of streams along this gradient. While canopy cover was not directly measured, abundant, dense filamentous algal streamers were observed at the NVR station. One of our 1^st^-order streams, WSB arises from soils with a high-water table, making the trees subject to wind-throw creating a more open forest canopy. The shift in community structure at the NVR site is consistent with decreased shading by riparian trees as stream order increases and follows predictions of the River Continuum Concept of shifts in the energy balance from allochthonous detrital carbon to autochthonous production in mid-order streams (16).

In this study, microeukaryotes contributed from 4.0 to 40.9% of microbial biomass and at the lower portion of the range (10% or less), PLFA profiles indicate heterotrophic microeukaryotes dominated while communities with larger proportions of microeukaryotes had a significant phototrophic component. This was evinced by the shift in the ratio of ω3 PLFA to ω6 PLFA. While ω3 and ω6 fatty acids are found in both heterotrophic and phototrophic microeukaryotes, they differ in their relative abundance (45-47). In heterotrophic eukaryotes, the abundance of ω6 and 3 fatty acids are comparable while in phototrophic eukaryotes ω6 fatty acids are typically found in abundances that are ~ 10% of the abundance of ω3 fatty acids. Across the streams in this study the relative concentrations ω6 and 3 fatty acids are consistent with phototrophic microeukaryotic comprising upwards of 75% of the microeukaryotic community of sediments with high percent microeukaryotes and the microeukaryotic community being dominated by heterotrophs in sediments with low percent microeukaryotes. The contribution of heterotrophic microeukaryotes to total microbial biomass in streambed sediments is likely limited by trophic interactions (unless seasonal and hydrodynamic conditions allow accumulation of significant particulate organic matter colonized by fungi), while the contribution of phototrophic microeukaryotes is likely limited by light availability and to a lesser extent biotic and abiotic disturbance and nutrient availability (48–50). Thus, it is clear that the presence of phototrophic microeukaryotes alters microbial community structure and function.

Our study showed that a critical, and often overlooked, factor in the ecology of stream sediments is the ability of open canopy/light (as seen in the NVR river) to shift the ratio of microeukaryotic to prokaryotic biomass and the nature of the reflected interactions. This is evinced by repeated finding that the major component of variation in sediment microbial community structure is driven by microeukaryotic/prokaryotic ratio as shown in this study (Fig. 3) and other studies (29, 44). For watersheds where shading of streams occurs, we posit the ratio of microeukaryotic to prokaryotic biomass is an essential component of any study striving to understand the critical roles sediment microorganisms play in the ecology and biogeochemistry of fluvial networks.

### Bacterial community structure

In this study, PLFA and DGGE analysis yielded similar patterns that showed distinct watershed differences in bacterial community structure (Figures 5a and 5b). The variation in bacterial community structure at regional scales may involve multiple causal pathways as a complex relationship between geology, bacterial community structure, and DOM quality exists. For instance, differences in watershed characteristics including bedrock geology and water chemistry generate differences in DOM qualities and quantities (40, 51), which in turn cause variation in stream sediment bacterial community structure (36). The complexity arises in that streambed microorganisms can, in turn, affect DOM quantity and quality (52). Wagner et al. (53) demonstrated clear shifts in bacterial community structure that included direct effects (increase relative abundance of cyanobacteria and other phototrophic bacterial taxa) as well as apparent indirect effects such as the increased relative abundance of predominately heterotrophic taxa such as *Roseomonas* and *Rivibacter* along with decreases in others such as *Planctomycetes* and *Gemmatimonadetes*. They also demonstrated, along with Rier et al. (54), a strong positive correlation among enzyme activities associated with heterotrophic bacteria and increasing light levels, and in the Rier et al. (54) study, these increases were greater than the increase in bacterial biomass.

Land use may be an alternative proximate cause for the observed variation in bacterial community structure between the two watersheds. The Neversink watershed is 95% forested while the WCC watershed, with its mixture of pastureland and forest, has a long history of agricultural land use, which is known to strongly affect sediment microbial communities (55). In addition, our data indicated significant differences in conductivity and sediment C:N mass ratios between the two watersheds, either of which can also significantly affect bacterial community structure. Microbial biogeography studies show that for geographic distance relevant to our study, deterministic processes primarily governed bacterial communities and their structure reflect local environmental heterogeneity, although distance effects are also noted (23-26, 56, 57). In this study, microbial community structure was influenced by light and total microbial biomass was strongly influenced by sediment carbon content, water content, C:N mass ratio, and surface area (see below). It is reasonable to assume that these factors may also influence bacterial community structure.

Besemer et al. (44) found greater alpha and beta bacterial diversity among headwater stream communities compared to larger streams of the same network. Although neither of the fingerprinting assays used in this study produce direct measures of diversity and the Besemer study (44) used biofilms removed from rock surfaces, our experimental design provides several tests of their hypothesis that headwater streams are a reservoir of bacterial diversity. These include a comparison of the variation in community structure found in the four 1^st^-order streams in the WCC watershed versus the 2^nd^ - and 3^rd^ - order WCC streams, and a comparison of the four 1^st^ - order Pigeon and Biscuit Brook tributaries to the 3^rd^ - order Pigeon and Biscuit Brooks. In three of these comparisons, sediments from the headwater streams showed substantially greater variation in bacterial community structure than sediments from the corresponding downstream stations while by one comparison (PLFA - Neversink streams) variation in bacterial community structure was greater among the downstream stations compared to the headwater streams (Figure 5a). These findings suggest that the observed pattern of decreasing bacterial diversity in stream biofilms with increasing stream order will likely extend to surficial streambed sediments. As dendritic ecological networks, fluvial systems experience a broad suite of coupled physical, chemical and biological processes that strongly influence microbial community structure (58). In the Eastern Deciduous biome, headwater streams are typically shaded and net heterotrophic, receiving appreciable inputs of terrestrial organic matter. Hierarchical patch dynamics within the local terrestrial environment, also resulting from a broad suite of coupled physical, chemical and biological processes, provide a varied terrestrial contribution to streamwater DOM, which in turn exert significant influence on bacterial community structure (36, 40, 44). While our study does not allow for direct measurement of alpha or beta diversity, the sampling density used for PLFA analysis does address aspects of a critical question within the riverscape concept, that is, the spatial scales of patch size for streambed sediments. Our results indicate that for these headwater streams there is a spatial scale of variation on the order of 10 cm. In addition, each reach contains multiple patches, a given patch (defined by microbial community structure - sensu (59)) can occur at multiple stations within a stream, and in multiple streams within the watershed (Figures 4). Using PC1 scores we calculated that the adjacent triplicate cores taken within a station encompassed anywhere from 10 to 100% of the variation observed for that stream. While this study did not directly investigate the impact of local heterogeneity on streambed microbial and bacterial community structure, the factors explaining variation in total microbial biomass likely contribute to variation observed in community structure.

### Microbial biomass and environmental variables

At the watershed and stream spatial scales, total microbial biomass showed significant differences but did not show a consistent trend within stations and triplet cores. Multiple regression analysis indicated percent sediment carbon content, percent water content, C:N mass ratio and sediment surface area explained nearly 70% of the variance in sediment biomass. Path analysis indicated that primary direct control was via sediment organic carbon, C:N ratios and sediment water content while sediment surface area affected sediment biomass indirectly via its impact on the other three variables. All four are known to be important environmental constraints of streambed microbial biomass and sediment organic carbon reflects a combination of several biogeochemical processes (60, 61) that influence microbial biomass through its quantity, quality or a combination thereof. Previous studies of stream sediments and terrestrial soils have shown the quantity of carbon significantly influenced microbial biomass (62-65). The results from this study further corroborate these observations and show that increasing sediment organic carbon concentration leads to greater total microbial biomass both directly and indirectly via interaction with C:N mass ratio and sediment water content (Fig. 8).

In addition, Findlay et al. (66) showed that variation in quality of sediment detritus, as measured by C:N mass ratio, was negatively correlated with bacterial abundance, while Schallenberg and Kalff (62) found variable results (either negative or no correlation) in lake sediments. Our results, that total microbial biomass correlated positively with both sediment organic carbon and C:N mass ratio suggests that organic carbon quantity can supersede the effects of quality, although increasing sediment C concentration combined with increasing C:N ratio might indicate an increased presence of fine particulate organic matter, which is known to be positively correlated with microbial biomass (67, 68). The cause of the difference between our findings and those of previous researchers is not known, however, the Findlay et al. (66) study did focus on particulate organic carbon; while in this study total sediment carbon was measured and the range of C:N mass ratios did differ in the two studies (9-27 vs. 6-17; (66) vs. this study, respectively).

Schallenberg and Kalff (62) working with lake sediments that ranged from 46% to 99% water content, found that percent water content was the single most important variable in predicting sediment bacterial biomass. They concluded that normalization of bacterial biomass to grams of sediment dry weight introduced a substantive bias, and recommended that microbial biomass be normalized to sediment surface area or sediment fresh weight. It is our experience that most studies are confined to sediments of a much narrower range of water content and that the usefulness of normalization to sediment dry weight (correcting for differences in sample size) far outweigh the risk of error introduction (36, 69). However, in this study sediments exhibited a wide range of sediment water content and required the recommended normalization to fresh weight. This is reflected in the finding that percent water content was the second most important variable in predicting sediment microbial biomass. High percent water content allows for greater aqueous connectivity within sediment, which in turn, allows nutrient and substrate transfer providing microorganisms with a continuous supply of nutrients, as well as, means to move to more favorable locations (70, 71). Nogaro et al. (72) working with sediment porosity (sediment water content corrected for adsorbed water) extends its importance to a deterministic process governing riverbed bacterial community.

Finally, sediment surface area has long been known to impact microbial communities, though many early studies (prior to the popularization of the Brunauer-Emmett-Teller method (73)) report this parameter in terms of sediment grain size (74, 75). Surface area of streambed sediments can influence microbial biomass through its effects on flow rates and availability of nutrients (76, 77) and quantity and quality of organic carbon (78, 79). In this study, path analysis indicated that the effect of sediment surface area on microbial biomass was indirect via its direct effects on sediment C:N mass ratio, and carbon and water content. A detailed study on how the surface area of the sediment affects the distribution of streambed microbiomes deserves further investigation.

## Conclusion

In conclusion, our study revealed regional-level patterns in microbial and bacterial community structures and adds to the growing number of studies suggesting that regional-scale environmental factors influence the biogeography of microbes. At the same time, we observed that local environmental factors strongly influence sediment microbial biomass, which can vary greatly among streams within a watershed, particularly among its 1st-order streams. Our findings highlight that variations in microbial and bacterial community structures within streams reflect a mosaic of small-scale patches and suggest that the type and spatial arrangement of patches is an important and often overlooked component of studies of metacommunity ecology of fluvial networks. Lastly, the fact that bacteria in headwater streams are typically components of a mixed-phylum community (80) and a growing literature that documents the impacts of microeukaryotes on microbial community structure, underscores a need to place research on bacterial community structure based on molecular methods into a broader context.

## MATERIAL & METHODS

### Study sites

Study streams were located within the 1^st^- 3^rd^ order, 7.3 km^2^ East Branch White Clay Creek (WCC) within the southern Pennsylvania Piedmont (39°53’N, 75°47’W) and the 1^st^- 5^th^ order, 171 km^2^ Neversink River in the Catskill Mountains of New York (41°57’N, 85°29’W) (Fig. 1). Predominant land uses within the WCC watershed upslope of an intact-forested riparian zone are row-crop agriculture (52%), hayed/grazed fields (22%), and wooded lands (23%) (81, 82). At the 3^rd^-order site, mean annual streamflow, stream water temperature, and local precipitation are 115 L/s, 10.6° C, and 105 cm, respectively (81). Soils are deep, unglaciated Utisols and streambed sediments consist of clay-, silt-, and sand-sized particles in pools and runs, with gneiss- and schist-derived gravel and cobble throughout all of the riffles; riffles from the western tributary and main stem of White Clay Creek also contain metamorphic materials derived from Cockeysville Marble. The dominant tree species are American beech (*Fagus grandifolia*), red oak (*Quercus rubra*), black oak (*Quercus velutina*), and tulip poplar (*Liriodendron tulipifera*) (81, 82). WCC flows into the Christina River, a tributary of the Delaware. The Neversink River watershed is contained within a mountainous region in northeast New York state with elevation ranging from 480 m to 1280 m. The hill slopes are steep with several deeply incised headwater channels and the soils in the Catskills region are predominantly acidic Inceptisols (83).

Streambed sediments consist of clay-, silt-, and sand-sized particles and shale-, siltstone-, sandstone- and conglomerate-derived gravel and cobble in riffles. At the 5^th^ -order site (USGS station 01435000), mean annual streamflow, stream water temperature, and local precipitation are 5.49 m^3^/s, 8.3° C, and 131 cm, respectively. The watershed is sparsely populated and 95% forested, primarily of mixed northern hardwood species dominated by American beech (*Fagus grandifolia*), sugar maple (*Acer saccharum*), and yellow birch (*Betula alleghaniensis*). Balsam fir (*Abies balsamea*) is common above 1,000-m elevation, and eastern hemlock (*Tsuga canadensis*) stands grow in a few areas that have poorly drained soils (83, 84). The Neversink River is a tributary of the East Branch Delaware River.

### Experimental design

We used a hierarchical design to evaluate spatial patterns of microbial biomass and community structure along a stream order gradient, where stream order refers to the Strahler (85) modification of the Horton (86) classification system. Our nested sampling design (Fig.1) focused on four spatial scales: 1) between watersheds (> 350km), 2) among streams within a watershed (50m-10km), 3) among sampling stations within a stream reach (2m - 25m), and 4) among cores within a sampling station (<1m). In WCC, we sampled one 3^rd^ - order reach adjacent to the Stroud Water Research Center in Avondale, Pennsylvania (WCC), the two 2^nd^ - order tributaries, White Clay Creek West (WCW) and White Clay Creek East (WCE), and four 1^st^ - order tributaries, two for each 2^nd^ - order stream, Ledyard Spring Brook (LSB), and Water Cress Spring (WCS) which flow into WCW, and Dirty Dog Spring (DDS) and Walton Spring Brook (WSB), which flow into WCE. The forest canopy varied between dense and open along the WSB bank. In the Neversink we sampled the 5^th^ - order Neversink River (NVR), two 3^rd^ - order tributaries of the West Branch Neversink River, Biscuit Brook (BBR) and Pigeon Brook (PBR), and four 1^st^ - order tributaries: Biscuit Brook Tributary A (BBA) and B (BBB), and Pigeon Brook Tributary A (PBA) and B (PBB). The United States Geological Survey maintains stream gauges at two of the Neversink sites (USGS 01434025, Biscuit Brook 300 m above the confluence with Pigeon Brook, and USGS 01435000, Neversink River near Claryville). Within each stream, three stations within a reach (downstream, midstream and upstream) were established and triplicate sediment cores were collected across each station as independent samples. In summary, the design consisted of 2 stream networks, 7 streams per stream network, 3 stations per stream and 3 replicate sediment cores per station, corresponding to a total of 126 samples of streambed sediments. Within the watersheds sampled, both rivers were unregulated. All streams within a watershed were sampled in the same week, and both watersheds were sampled within a 2-week period in July and August 2010 to reduce seasonal differences.

### Sampling procedures

Samples were delimited with a 100mm diameter plastic ring that was inserted 2cm-deep into the streambed (a 75mm diameter ring was used for 1st - order streams whenever streambeds were dominated by large rocks, cobbles or stones). Plexiglas© plates were slipped under and over the ring to effectively trap the sediments and allow them to be lifted intact from the streams with minimal disturbance. Sediments in the top 2mm within the ring were transferred with a sterile spatula to pre-labeled Whirl-Pak sampling bags and stored on ice prior to subsampling. Stream water conductivity and water temperature readings were measured with a YSI model 32 conductance meter. Within six hours of sampling, sediments were transferred to a clean plastic weigh boat, thoroughly homogenized, subsampled for phospholipids, DNA, surface area, or elemental analyses, and frozen. Frozen samples were shipped to the appropriate laboratory for analysis.

### Sediment surface area, particle size and elemental analyses

Frozen subsamples for surface area and particle size analyses were dried at 60°C and split with one subsample being processed through a U.S. Standard sieve series (W. S. Tyler Co., Menton, Ohio) for sediment particle size distributions. The other subsample was heated to 350°C to remove organic matter and analyzed by the Brunauer–Emmett–Teller (three-point adsorption isotherm) method using a Micromeritics Tristar surface area and porosity analyzer (Micromeritics Corporation, Norcross, GA) and N_2_ as the adsorbate (73).

The frozen subsamples for elemental analysis were freeze-dried, finely ground, and inorganic carbonate removed (gaseous HCl). Approximately 35 mg of sediment were analyzed on a Costech 4010 elemental analyzer for percent carbon and nitrogen, and atomic carbon to nitrogen ratio (C:N).

### Phospholipid analysis

Microbial biomass and community structure were determined using phospholipid phosphate (PLP) and phospholipid fatty acid (PLFA) analyses following the methods of Findlay (45). Briefly, cellular lipids were extracted from the frozen sediment samples by dichloromethane/methanol/water extraction and partitioned into aqueous and organic fractions. The organic fraction containing the lipids was subsampled for PLP analysis (87). PLFAs were recovered from other lipids by differential elution from silicic acid columns (J. T. Baker, Center Valley, PA, USA) and were analyzed as their methyl esters. Purified fatty acid methyl esters (FAME) were identified and quantified using gas chromatography. The FAME were analyzed by gas chromatography in an Agilent gas chromatograph equipped with an automatic sampler, a 60 m x 0.25 mm non-polar DB-1 column, and a flame ionization detector. Hydrogen was used as the carrier gas at a flow rate of 2.3 ml/min. Oven temperature was 80 °C at injection, increased at 4 °C/min to 250 °C and held at 250 °C for 10 min. FAME identification was based on relative retention times, coelution with standards, and mass spectral analysis. The FAME nomenclature used followed Findlay and Dobbs (88). Using polyenoic fatty acids as indicators of microeukaryotes, total microbial biomass was partitioned between prokaryotic and microeukaryotic organisms and the results were presented as percentages (88).

### Bacterial community structure analyses by PCR-DGGE

Genomic DNA was extracted from 0.3 g subsamples of the frozen subsamples using the Power Soil DNA Extraction kit (MoBio Laboratories, Carlsbad, CA, USA) following the instructions from the manufacturer. DNA was quantified by spectrophotometric absorption at 260 nm, and the purity was assessed from absorbance ratios at 260/280 and 260/230 nm using a ND-2000 Nanodrop spectrometer (Thermal Scientific, Wilmington, DE). 16S rRNA genes were amplified with 1070f (ATGGCTGTCGTCAGCT) and GC-clamped 1392r (ACGGGCGGTGTGTAC) primers (89). The PCR reactions were performed using an automated Eppendorf Mastercycler Thermal Cycler (PerkineElmer, Norwalk, CT). PCR-DGGE was performed using the Dcode system (Universal Mutation Detection System, Biorad) as previously described (89, 90). Briefly, equal amounts of PCR products were loaded onto an 8% vertical polyacrylamide gel containing a 50-70% denaturing gradient made of urea and formamide. Gels were electrophoresed at 60 °C and 70 V for 16 h and visualized with SYBR Gold staining (Life Technologies, NY). Digital photographs of gel images were analyzed using GelComparII v.5.10 (Applied Maths, Austin, TX, USA) and visually checked for accuracy. Distinguishable bands represented distinct bacterial taxa that occurred in each sample (89, 90), and these data were used for community structure and downstream multivariate statistical analyses (below).

### Statistical analyses

Nested analysis of variance (stations nested within streams, and streams within watersheds) with Turkey’s HSD (p < 0.05) was performed on sediment organic content and microbial biomass to determine significant differences across spatial scales (JMP 10 and MINITAB 16). Across streams, sediments varied greatly in their percent water content, violating the assumption necessary for standardizing data to sediment dry weight (62). Hence, we reported biomass and abundance per gram of fresh weight sediment, rather than the customary dry weight of sediment. Relationships among variables were investigated using linear regression and multiple regression analyses (MINITAB 16). We tested data for normality with the Shapiro-Wilk test and homogeneity of variance with the Bartlett test with appropriate transformations applied as needed. For multiple linear regression analysis, predictor variables (environmental variables) were selected using the ‘best subsets’ algorithm in MINITAB. This algorithm fits a small fraction of all possible regression models and reports the ‘best subset’. We identified the best model based on several selection criteria including adjusted r^2^ and Mallows Cp (91). We used path analysis, a specific form of structural equations modeling (SEM), to explore further the direct and indirect influence of environmental variables (as independent variables) on microbial biomass (as dependent variable) using the SAS Structural Equation Modeling subroutines for JMP 10. SEM is a multivariate statistical technique that tests the importance of pathways in hypothesized models, and allows comparison of models to experimental data (92). Standardized regression coefficients between variables were calculated and plotted as path coefficients on path diagrams constructed for microbial biomass. Natural log transformed (ln (wt% + 1)) PLFA relative abundance data were subjected to principal component analysis (PCA) to identify patterns of variation in the microbial community structure across spatial scales and stream orders. PCA was performed for the combined data set of Neversink and WCC networks (SPSS 19). PLFA profiles were interpreted using a functional group approach (88). DGGE data were entered as presence or absence of bands, pairwise comparisons calculated and bacterial community structure analyzed by NMDS (Nonmetric Multiple Dimensional Scaling) employing the MDS procedure in SAS/STAT Software (v 9.3, SAS Institute Inc., Cary NC). The bacterial community distributions were illustrated in 2-D NMDS plots.

## Acknowledgments

Sherman Roberts, Michael Gentile (Stroud Water Research Center, Avondale, PA) and Janna Brown (University of Alabama, Tuscaloosa, AL) assisted in sample collection and processing. Christina Staudhammer provided invaluable advice on the application of path analysis; however, the authors take full responsibility for the application and interpretation of all statistical analyses. Funding for this project was provided by the National Science Foundation grants to RHF (DEB-0516235 and DEB-1119922) and LAK (DEB-0516449 and DEB-1120717).

